# Conserved gene- and network-level thermal memory intervals in two divergent perennial crucifers in nature

**DOI:** 10.1101/2025.10.31.685746

**Authors:** Yoshikazu Endo, Haruki Nishio, Taichi Oguchi, Kyoko Yamane, Victoria Faith Eseese, Clarissa Frances Frederica, Hiroshi Kudoh, Diana Mihaela Buzas

## Abstract

Some biological responses persist long after the initial stimulus has disappeared—a phenomenon known as cellular memory. In its long-term form, cellular memory often reflects interactions between *cis*-acting chromatin states and diffusible *trans*-acting regulators, which are experimentally difficult to separate *in vivo*. A key challenge is to develop a quantitative framework that captures cellular memory length even without prior mechanistic knowledge. The *FLOWERING LOCUS C* (*FLC*) gene illustrates this problem and opportunity: a Polycomb/Trithorax *cis*-acting chromatin switch at *FLC* produces bistable ON/OFF transcriptional states, while *trans*-acting factors such as *VIN3* and *FT* modulate transitions between those states. In laboratory studies, *FLC* memory is classically viewed as persistence of repression or activation after the inducing signal disappears; field studies additionally show that *FLC* integrates fluctuating temperatures over past intervals, revealing time-integrative mode of memory. To quantify such long-term effects systematically, we formalized the thermal memory interval (TMI), the time window of past environmental cues that best predicts current gene expression—as a transferable metric. We applied TMI to the *VIN3–FLC–FT* module in two perennial *Brassicaceae* with divergent life histories: *Arabidopsis halleri* subsp. *gemmifera*, an established ecological model, and *Eutrema japonicum*, introduced here to identify features generalizable beyond a single species. Using long-term expression and meteorological data, TMIs distinguished spring versus autumn *FLC* states and revealed distributed memory across the VIN3-FLC-FT network, with intervals from 1–150 days, extending previously reported FLC timescales. While TMIs require dense time-series data and do not by themselves reveal molecular mechanism, they offer a robust, quantitative, and generalizable framework: TMIs can be extended to other genes and to alternative environmental or physiological variables, enabling direct, comparative quantification of cellular memory across genes, species, and contexts.

## Introduction

Cells have the capacity to retain information about past stimuli, a phenomenon broadly referred to as cellular memory. Such memory enables biological systems to mount appropriate responses long after the initiating signal has disappeared. In animals, immunological memory provides lasting protection against previously encountered pathogens (1). Developmental systems also rely on cellular memory; for instance, in *Drosophila*, the specification of body segment identity during embryogenesis is stably maintained throughout the organism’s life. In plants, exposure to prolonged winter cold leads to vernalization, a classic example of cellular memory that ensures flowering occurs only under favourable spring conditions (2). Mechanistically, cellular memory, especially on long term scales, can be sustained through self-reinforcing genetic feedback loops or chromatin-based mechanisms, which often act together in complex ways that are difficult to disentangle (3).

A wide range of chromatin regulators, including Polycomb and Trithorax Group (PcG/TrxG) proteins, stabilize gene expression in bistable ON and OFF states and allow these transcriptional states to be maintained long term across cell divisions (3). At molecular level, variation in duration of cellular memory arises from differences in how readily these states can be reversed—through both gene silencing and reactivation—which depend on the intrinsic kinetics of the chromatin regulator, the organization of gene networks at native loci and environmental/developmental contexts. As a result, cellular memory varies across loci, environmental conditions, and species (4). While synthetic single-cell assays have capacity to measure and compare cellular memory lengh amongst different chromatin regulators, an experimental limitation is their observation windows, typically spanning only hours to days. At the opposite extreme, developmental and physiological readouts reveal that signals can be sustained over timescales of weeks, years or even across generations (1,2,5,6) but such outcomes are indirect and system specific. Taken together, the field currently lacks a general methodological framework for quantifying the duration of long-term cellular memory to allow comparisons across loci, contexts, and organisms.

Vernalization has become a powerful model to study long-term cellular memory through studies in both controlled laboratory assays and natural populations. From laboratory assays, vernalization is classically understood as the persistence of a response after the initiating signal disappears. The central vernalization integrator *FLOWERING LOCUS C* (*FLC*), a floral repressor, exemplifies this principle: its active and silenced states are initiated by developmental and environmental cues, respectively. Once cold is removed experimentally, cellular memory is evident at both molecular and physiological levels—through sustained *FLC* repression and the promotion of flowering (2). Conversely, when the embryo-specific transcription factors that activate *FLC* are no longer present, the active *FLC* state persists until plants experience prolonged cold, maintaining vegetative growth (7).

Beyond serving as a model for physiological memory, rare integration of experimental and mathematical approaches (8–10) in *Arabidopsis thaliana* also provided one of the first mechanistic demonstrations that PcG/TrxG-modified chromatin can encode memory through bistable ON/OFF states *in cis* (8–11). While at single-cell level *FLC* is transcriptionally ON or OFF, at the tissue level these states combine into four quadrants with distinct chromatin signatures and biological functions (12). The “silenced” and “active” *FLC* expression states are maintenance states and correspond to all cells being transcriptionally OFF or ON, referred to here as the *FLC* Minimum and *FLC* Maximum states, respectively. The intermediate phases, where mixed ON/OFF cell populations cause gradual shifts in total transcript levels are “dialling” phases which arise from reversible switching between ON and OFF transcriptional states. During *FLC* dial-down, progressive switching to the OFF state allows the degree of silencing to adjust to the duration of cold (13); during *FLC* dial-up, switching to the ON state reinstates high *FLC* expression early in development, resetting the vernalization requirement and preventing premature flowering (14,15). This combination of reversible switching and long-term bistability shows that FLC behaves as both a stably maintained and dynamically regulated gene—an uncommon property among PcG/TrxG targets, which are typically classified as one or the other (Reinig et al., 2020). Therefore, the discovery of a *cis*-acting chromatin switch at FLC provided the mechanistic basis of time measuring during winter and expands the classical view of vernalization memory beyond persistence after disappearance and reveals a combination of features otherwise found in distinct classes of PcG/TrxG-regulated genes (16,17).

A series of studies extended vernalization research from controlled laboratory assays to natural field conditions and from annual to perennial life histories (18–21). Among these, Aikawa et al. (2010) identified a distinct form of *FLC*-based memory, defined not by persistence after signal disappearance but by the integration of fluctuating temperature signals over time. The *cis-*acting chromatin switch characterized in annuals (10,22) also operates in perennials (21,23), acting together with *trans*-acting factors such as the orthologues of *VERNALIZATION INSENSITIVE 3* (*VIN3*; (24)) and *FLOWERING LOCUS T* (*FT*; (25)), both showing seasonal expression patterns (20).

Aikawa et al. (2010) provided three key insights. First, the evergreen perennial *Arabidopsis halleri subsp. gemmifera* (*A. halleri*, hereafter, and *Ahg* as a prefix of gene name) allows continuous monitoring of *FLC* cellular memory throughout the annual cycle in dividing leaf cells, revealing a cyclic reiteration of *FLC* maintenance and dialling states across seasons. Second, despite multiple environmental cues influencing flowering in perennials (26), temperature alone explained ∼80% of *AhgFLC* expression variance when averaged over a six-week window, demonstrating that *FLC* integrates temperature across time. The strong predictive power of a single factor highlights an exceptional form of cellular memory: integration of a fluctuating environmental signal over a defined temporal window. Third, Aikawa et al. (2010) introduced an empirical approach to capture this property by regressing gene expression against past temperature windows. Building on this framework, Nishio et al. (2020) incorporated bootstrap analyses to identify robust best-fit intervals. Here, we formalize this approach as a transferable metric—the Thermal Memory Interval (TMI)—which quantifies the timescale over which past temperature history predicts current transcriptional states.

Rather than seeking to dissect molecular mechanisms or define functional roles of individual loci, this study leverages the experimental advantages of natural populations of evergreen perennials growing in their native environments to evaluate whether TMI represents a quantifiable property at both gene and network scales. Specifically, we asked whether TMIs can be measured robustly and whether they provide biologically informative insights into chromatin stability, transcriptional responsiveness, and delayed gene expression over seasonal timescales. To establish this foundation, we first characterized vernalization in a natural population of the divergent perennial relative of *A.halleri*, *Eutrema japonicum* (*E.japonicum*, *Ej* as a prefix of gene name). Comparative analysis is informative because unlike the atypical life history of *A.halleri*, where all meristems transition to reproductive fate before reverting to vegetative growth, *E. japonicum* exemplifies a more typical perennial strategy, with only some meristems becoming reproductive during winter while others remain vegetative to sustain growth into the next cycle. We first confirmed that the *FLC* orthologues in *E. japonicum* represses flowering and that prolonged cold exposure relieves this repression. To this end, we pursued three complementary approaches. First, we analyzed high-resolution gene expression data with environmental temperature records, adapting methodologies from (18) and (21) to calculate TMIs and assess their predictive power. Second, we compared TMIs across the two divergent perennials, applying them as general ecological parameters to classify cellular memory types and identify conserved features. Analysis of these data, together with chromatin and transcriptional profiling of *EjFLC* in autumn and spring, demonstrated that TMIs reliably differentiate *FLC* dialling states and provide predictive insight beyond direct molecular measurements. Finally, transfer experiments tracking the *EjVIN3*–*EjFLC*–*EjFT* network dynamics under non-native but natural environments revealed robust network configuration, while also showing that temperature variation experienced up to six months prior left a detectable mark. Notably, the thermal memory interval of *FT* exceeds that of *FLC*, a feature conserved across divergent perennial life histories, extending the known range of thermal memory from six to 21 weeks and suggesting cumulative information flow within the VIN3– FLC–FT network. Together, these approaches position TMIs as a tractable ecological parameter for quantifying cellular memory across genes, networks, and species.

## Materials and Methods

### Study sites and phenotyping

We selected a natural population of *E. japonicum* situated in the Ikawa Experimental Forest of the University of Tsukuba Mountain Centre, in Shizuoka, Japan at 35°13’ N, 138°14’ E, 710 m in altitude, where plants grow in a flat, shaded area with water running down during restricted times of the year. For *A.halleri*, the natural population location was described (27). For the September and October Tsukuba transfer experiments, *E. japonicum* plants were placed outside in Tsukuba, Ibaraki at 36°05′ N, 140°14′ E, 26 m altitude. *E. japonicum* plants scored as “flowering” included visible flower buds and all stages of bolting. A minimum of 12 random plants with similar root diameter (about 1-1.5cm) was included in each sample.

### Plant and tissue sampling

For *E. japonicum*, a repository of all samples is included in supp Table S1. About 200-300 mg leaf tissue was pooled from at least six different plants for each of the three biological replicates and immediately placed on dry ice (Ikawa series 2016-8) or at -80**°**C (Tsukuba series 2018-2019), except for the 2021-2022 series, where plants (subterranean stem portion and all aerial parts) were first placed at 4**°**C for a maximum of 24 h during transport to the laboratory, before freezing at-80°C. For the 2021-2022 series, *E. japonicum* stems of more than 1 cm in diameter were detached in Ikawa from 12 plants per treatment, transported to the laboratory, and planted into cactus soil mix pots (S8 Fig); the first sampling was performed when new growth was confirmed approximately four weeks after transfer. Sampling was done during 12.00-13.00 (Tsukuba) and 14.00-16.00 (Ikawa). For temperature transfers, two sets of re-potted plants for each “winter” (December 19th, 2018) and “spring” (March 19th, 2019) were placed into growth chambers at 4**°**C or 23**°**C and further divided into 14- and 28-days subsets. Sampling in *A.halleri* was performed as previously described (19,21).

### Transgenic *A. thaliana* assays

cDNA prepared from *E. japonicum* RNA sampled on September 21, 2016, was amplified with attb1 and attb2 oligos using PrimeSTAR HD DNA polymerase (Takara, R010A). The products were recombined into pDONR221 (Invitrogen, 12536017) using the Gateway BP Clonase II enzyme mix (Invitrogen, 11789-20) and 12 clones were sequenced. One *EjFLC1* and *EjFLC2* clone was recombined into the pB2GW7 binary vector (28). The final plasmids were transformed into *Agrobacterium tumefaciens* strain *GV3101* and transformed into the *flc3* mutant in the Col-0 background using floral dip (29). Flowering was scored as days to flower or DNF (did not flower), when no flower emerged in the first 100 days from planting, in 50 control and transgenic T0 transformants for each construct.

### Genomic and cDNA sequencing, RNA extraction, quantitative RT-PCR and Chromatin Immunoprecipitation

Genomic sequences were first identified from the commercial *E. japonicum* ecotype Mazuma No. 3 (30) based on homology with queries against *AtFLC*, *AtVIN3*, and *AtPP2A3*, while *EjFT* sequence was known (31).We amplified genomic FLC PCR fragments spanning 5.9 kb from ATG to approximately 1.3 kb downstream of the stop codon from an individual plant from the Ikawa ecotype and fully sequenced 12 clones at Eurofins Genomics. For *E. japonicum*, RNA was extracted using RNeasy Plant Mini Kit (Qiagen, Hilden, Germany), and cDNA was prepared from 2 μg of RNA using Superscript III (Invitrogen 18080051). Q-RT PCR and ChIP q-PCR were performed on three biological replicates, each with three PCR repeats, and carried out using Toyobo THUNDERBIRD SYBR qPCR Mix (QPS-201) and ABI7900. Expression levels were calculated using the ΔΔCT method (32), and for the ChIP experiment, the absolute DNA amount was quantified using input DNA for the standard curve. ChIP amplicons were designed for regions with 100% sequence identity between *EjFLC1* and *EjFLC2*. For *A.halleri,* all procedures were performed as previously described (21). All primers are listed in Table S2.

### Statistical modelling

Temperature records were obtained from the nearest Japan Meteorological Agency station (35°13′ N, 138°13.3′ E 755m altitude in Ikawa, Shizuoka; 36°6.2′ N, 140°13.2′ E, 26m in Tsukuba, Ibaraki for *E. japonicum* and 34°59.9′ N, 134°59.8′ E, altitude 72m, Nishiwaki, Hyogo Prefecture, Japan for *A. halleri*). Simple moving averages (SMA) of temperatures were calculated for the time windows of the previous 1 d − 150 d from each sampling date. Gene expression was modelled using the SMA of temperature for each time window by linear regression using the lm function in R v4.1.1. The R2 value was calculated for the SMA of the temperature for each time window. One thousand bootstrap data sets of gene expression and SMA of temperature were created to estimate the median and 95 % intervals of the R2 values. For all bootstrap datasets, we calculated the median and 95 % intervals of the best SMA period, which explained the gene expression at the highest R2 value (Fig 6).

The intervals used to calculate the SMA were from two seasons, as follows: for *E. japonicum*, September 7, 2016, to February 22, 2017, for *EjFLCs* dial-down and March 8, 2017, to May 31, 2017, and from March 14, 2018, to May 23, 2018, for *EjFLCs* dial-up; for *A.halleri* November 6, 2012, to February 19, 2013, and from November 5, 2013, to February 18, 2014, for *AhgFLC* dial-down and March 7, 2013, to May 28, 2013, and from March 4, 2014, to May 27, 2014, for *AhgFLC* dial-up.

## Results and discussion

### The divergent perennial life histories of *Arabidopsis halleri* subsp. *gemmifera* and *Eutrema japonicum* provide distinct experimental opportunities

Tracking long-term cellular memory depends on the ability to follow gene expression ideally in the same cell lineage and over natural timescales. Here, we focus on two perennial *Brassicaceae* with divergent life histories, *A.halleri* and *E. japonicum* (Fig 1A,B), each offering distinct experimental opportunities. *AhgFLC* is already well characterized in natural populations (18,20,21,33), however, meristems in *A.halleri* are atypically indeterminate: all vegetative meristems transition to reproductive growth after winter and then revert to vegetative fate in spring, producing aerial rosettes for the next cycle (Fig 1, S1 Fig). This unusual meristem biology is expected to be tightly linked with the kinetics of *AhgFLC* dial-up (18,33) and, by extension, with the dynamics of all four FLC regulatory phases (12). To broaden this framework, we introduced *E. japonicum*, also an evergreen perennial *Brassicaceae* relative but so far little explored in vernalization studies. *E. japonicum* represents a more typical perennial system in that it has determinate meristems: vernalization triggers flowering in only a subset of meristems, while others remain vegetative to sustain growth into the next cycle. Moreover, in *E. japonicum* the meristems giving rise to reproductive growth are located in the upper tiers and are therefore spatially distinct from vegetative meristems (Fig 1A). This clear distinction between rosette and cauline leaves, present in *E. japonicum*, disappears in *A.halleri* towards the end of spring. *A.halleri* produces primarily cauline leaves in April and May, limiting rosette leaf availability for sampling during this period (19,21). Therefore, *E. japonicum* permits continuous, year-round sampling of one lineage, i.e dividing rosette leaves—a feature not possible in *A.halleri*—providing a more direct route to monitoring the four regulatory phases of FLC across a full annual cycle. By comparing the two species, it becomes possible to distinguish more general properties of long-term cellular memory from aspects of FLC regulation that depend on meristem biology.

**Fig 1.**
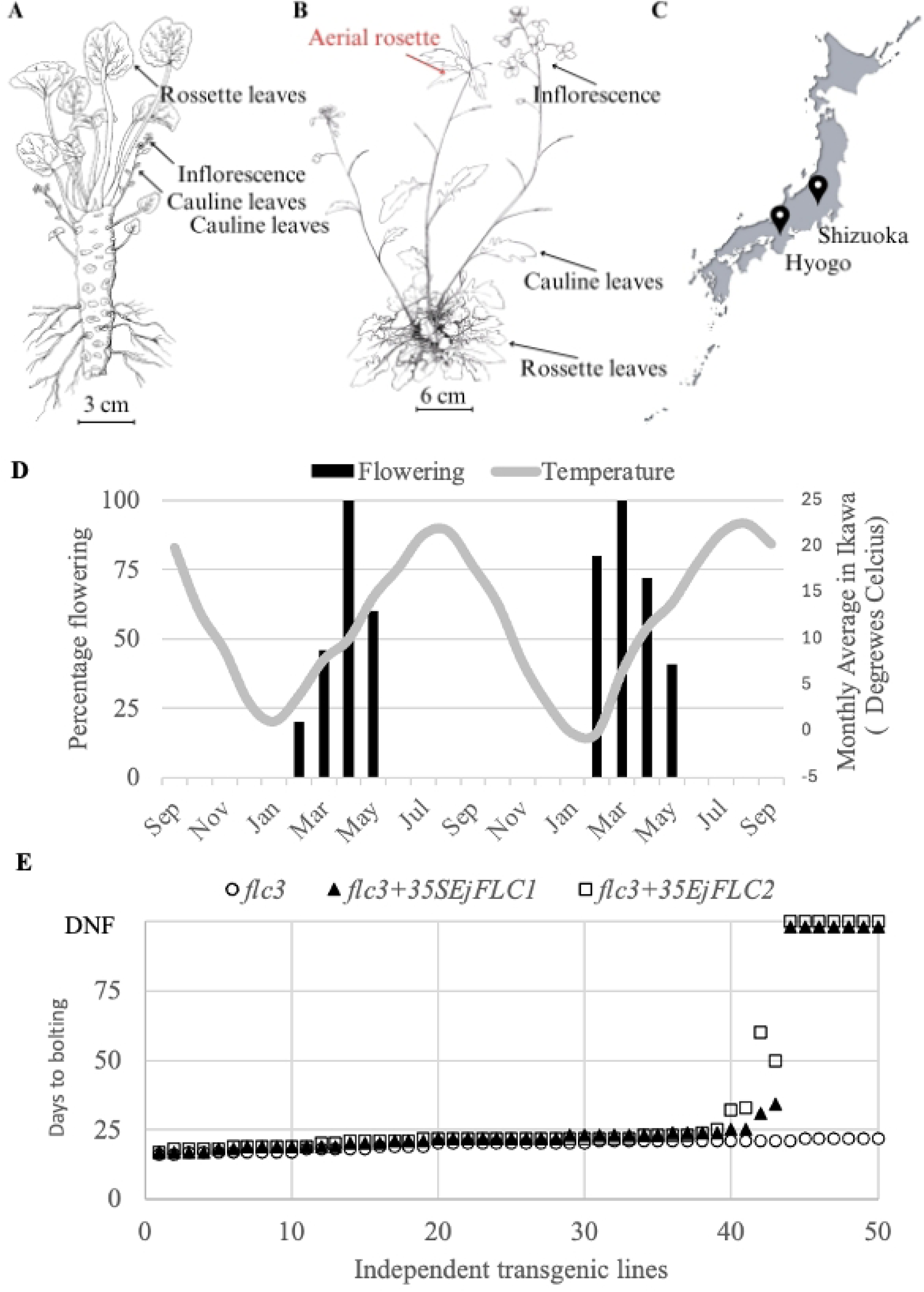
Perennial life history in two divergent *Brassicaceae* perennial life hisotires, *E. japonicum* vernalization response and transgenic assay in *Arabidopsis thaliana*. (A) General *E. japonicum japonica* biology (re-drawn from “Wasabi no subete”, page 9). Cauline leaves, present on inflorescence steams, are easy to distinguish spatially from rosette leaves. (B) General *Arabidopsis halleri* subspecies *gemmifera* biology. All meristems formed before and after the cold season develop into inflorescence stems and all inflorescences revert into vegetative growth and eventually form aerial rosettes (also see S1 Fig). Note that three types of floral stems are drawn in representative stages on the same plant, yet this configuration cannot be found in natural settings at the same time. (C) Locations of the natural population of *Eutrema japonicum* in Ikawa, Shizuka and *Arabidospis halleri* subspecies *gemmifera* in Nishiwaki, Hyogo. (D) *Eutrema japonicum* flowering interval plotted against the monthly average temperature in Ikawa. (E) Complementation assay with two *Eutrema japonicum FLC* genes (*EjFLC1* and *EjFLC2*) in the *Arabidopsis thalian*a *flc3*Col-0, estimated by flowering time in 50 independent transgenics, plotted in ascending order of days to flower, ending with plants which Did Not Flower (“DNF) during the 100 days of the experiment.

### Flowering in a *E. japonicum* natural population is synchronized to early spring and it is inhibited by *FLC* orthologues

To investigate whether *E. japonicum* exhibits a vernalization response under natural conditions, we first determined the flowering interval in a selected natural population of *E. japonicum* from Ikawa, Shizuoka (Fig 1C). In agreement with previous studies (31), differentiated flower buds became visible in February, the peak of flowering was in March and flowering ended by early May; also, plants remained vegetative outside this interval of the year(Fig 1D). Therefore, floral buds were initiated under the short days immediately following the coolest time of the year, in two consecutive years (Fig 1D). The finding that in *E. japonicum* prolonged low temperatures, but not long days, are critical for flower initiation is consistent with other perennial crucifers (34,35), including *A.halleri* (*18,21*). Therefore, *E. japonicum* undergoes vernalization in natural environment to synchronize flowering to late winter/early spring.

To characterize the *FLC* orthologues in *E. japonicum*, we first sequenced cDNA clones spanning the start and stop codons of *EjFLC* sampled throughout the year, from the Ikawa ecotype. We found six single nucleotide polymorphisms resulting in two amino acid differences, likely encoding two FLC proteins, and we termed the genes *EjFLC1* and *EjFLC2* (S2A Fig). To test if *EjFLCs* can inhibit flowering in the reference *A. thaliana*, complementation tests were performed with each gene in loss-of-function *A. thaliana flc3*Col-0 mutant. Flowering was delayed to a different extent in most primary transgenics of both *flc3*Col-0 *35S:EjFLC1* and *35S:EjFLC2* comparing to the *flc3*Col-0 and furthermore, 14% of *flc3*col *35S:EjFLC1* and 18% of *35S:EjFLC2* lines did not flower during 100 days in our experiments (Fig 1E). This data indicated that both *EjFLC1* and *EjFLC2* are similarly potent floral inhibitors, as expected form high amino acid identity between *AtFLC* and *EjFLCs* (S2B Fig). The high sequence similarity between *EjFLC1* and *EjFLC2* prevented us from quantifying individual *EjFLC* mRNAs. To understand if *EjFLC* genes are differently regulated throughout the year, we determined how *EjFLC1* and *EjFLC2* are represented in cDNAs prepared from samples from six months (S3 Fig). As we did not detect any major bias of one gene or the other we performed quantitative analysis of the sum, labelled “*EjFLCs”* from here onwards.

### Two divergent *Brassicaceae* perennial life histories mirror multiple gene expression characteristics

The four-quadrant *FLC* expression cycle in *A.halleri* reflects a temperature memory system year-round (18). To test whether this pattern represents a broader feature of *Brassicaceae* perennials, we conducted a high-resolution, two-year expression analysis of the three core vernalization pathway orthologues genes, *VIN3*, *FLC*, *FT* in the natural populations of *E. japonicum* from Ikawa, Shizuoka against the previously studied *A.halleri* natural population from Hyogo (20,21) (Fig 1C).

Despite having divergent life histories, both species displayed striking conservation of multiple expression features. First, both *AhgFLC* and *EjFLC*s expression levels spanned up to over 10^3^-fold under natural field conditions. By spring, steady-state levels dropped by approximately 10^3^fold (Fig 2 B and E). This large dynamic range in the field far exceeds the ∼10-fold change typically observed after saturating vernalization intervals in laboratory experiments, regardless of sample type or life history (36,37). Indeed, in *A.thaliana* the reduction in *AtFLC* levels under field conditions in a winter annual ecotype (38) was also in of a greater magnitude than under laboratory conditions, indicating that a large amplitude of *FLC* expression under natural conditions is conserved across annual and perennial life histories. These results suggest considerable differences in transcription rate and or mRNA degradation rate may occur between laboratory and field conditions. Alternatively, consistent with ON/OFF transcriptional states, it could be that *FLC* becomes activated in a larger fraction of dividing cells of leaves under natural conditions. This elevated transcription level would facilitate the observed graded response over the long-time plants encounter low temperatures in autumn and winter (Fig 2B, September to March; 2E, November to March).

**Fig 2.**
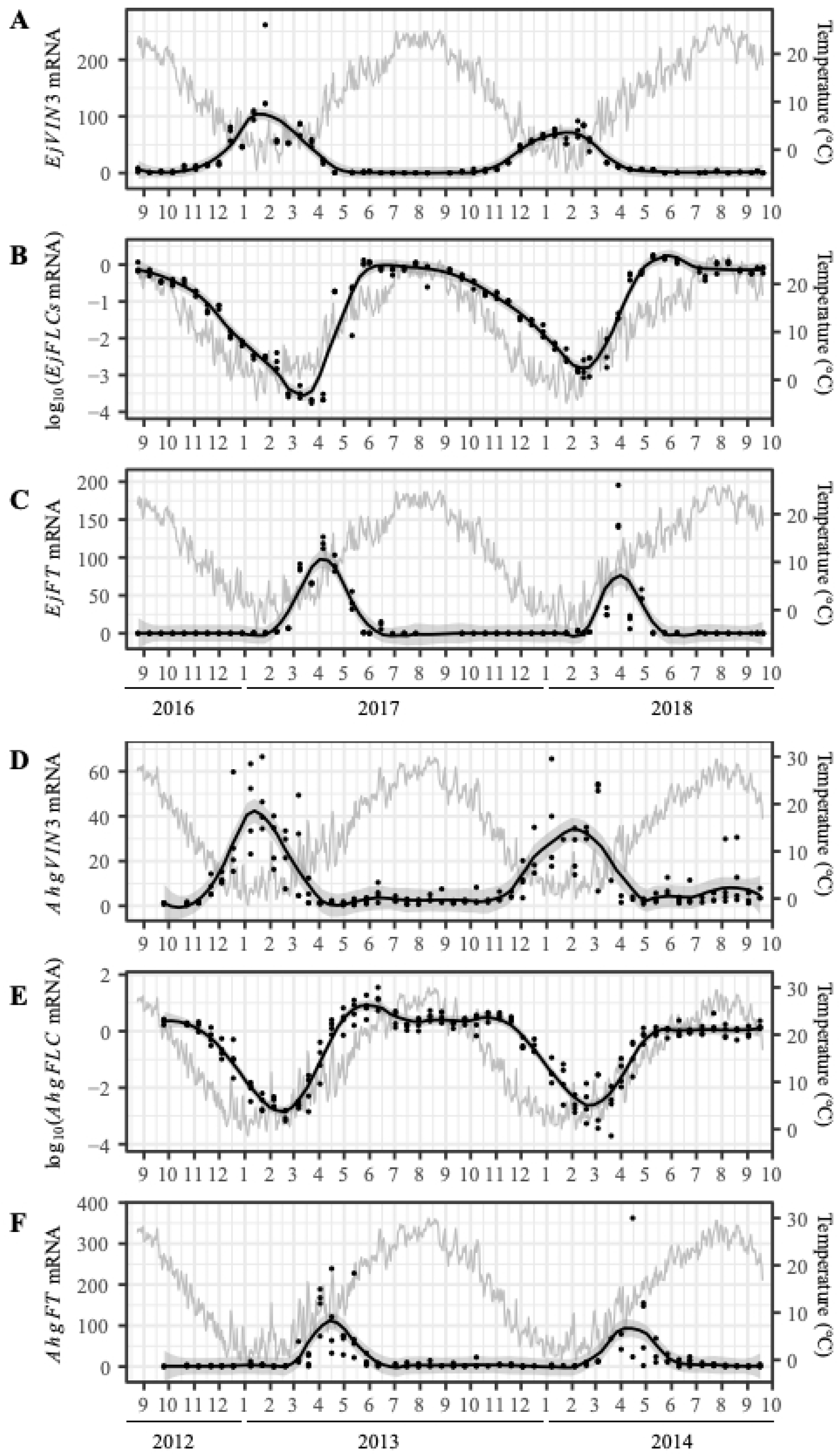
Relative quantification of *VIN3*, *FLC*, and *FT* mRNA in natural populations of two divergent perennials over two consecutive years. Average of three biological replicates of mRNA quantification for all genes, as indicated on the vertical axis, using ΔΔCT method. Internal controls were *EjPP2A3* in *Eutrema japonicum* (A-C) and *AhgACT2* in *Arabidopsis halleri subspecies gemmifera* (D-F).

Second, the spline curves of *AhgFLC* and *EjFLCs* gene expression are tilted, revealing two asymmetries: between the maintenance quadrants (*FLC* maximum and *FLC* minimum) and between the dialling quadrants (*FLC* dial up and *FLC* dial down) (Fig 2 B,E). We observed that *FLC* expression remained at a minimum for approximately 5 weeks in *E. japonicum* and 6 weeks in *A.halleri*. These durations are considerably shorter than the maximum *FLC* expression phases, which lasted about 14 weeks in *E. japonicum* and 22 weeks in *A.halleri*. These timelines also contrast sharply with controlled vernalization assays, which typically show an FLC minimum lasting for example about 7 days in the *Arabis alpina* perennial (39) and in *A.halleri* (40). These results suggest that, counterintuitively, fluctuating environmental conditions in nature may promote longer term stability of *FLC* maintenance states compared to the constant conditions of laboratory settings.

Surprisingly, *FLC* dialling states are asymmetrical between autumn and spring, even though seasonal temperature ranges are comparable (Fig 2B,E). In *E. japonicum*, *EjFLCs* dial-down lasted 18 weeks, whereas dial-up took only 9 weeks. A similar bias was observed in *A.halleri* (12 vs. 6 weeks). Previous vernalization assays in both annual and perennial plants suggest a similar tendency to asymmetry in *AtFLC*, *AaFLC*, *AhgFLC* expression (14,21,36,41) but such comparisons are complicated by measurements across different cell lineages. For example, the *AtFLC* dial-down takes place in leaves while *AtFLC* dial-up in embryos (42). Our *E. japonicum* analysis under natural conditions overcomes this limitation by tracking both processes in the same cell lineage, namely rosette leaves. These results provide the first direct evidence that *FLC* dial-up and dial-down are not simply equivalent mirror-image all-or-none activation and silencing events but instead display a seasonal bias. This supports the view that *trans*-acting inputs modulate the *cis*-encoded *FLC* chromatin switch throughout the year (23).

Third, annual expression patterns of *VIN3*, *FLC*, and *FT* formed a consistent regulatory sequence across species. In both *A.halleri* and *E. japonicum*, *VIN3* peaked before *FLC* Minimum, and *FT* peaked immediately afterward (Fig 2). This VIN3–FLC–FT domino-like effect unfolds in slow motion over 5–6 months in both *E. japonicum* (Fig 2A-C) and *A.halleri* (Fig 2 D-F), revealing features that are difficult to capture in short laboratory assays. First, *FLC* began to decline before *VIN3* expression peaked—late October in *E. japonicum* and late November in *A.halleri*—suggesting that an early, VIN3-independent phase of *FLC* repression, supported by genetic studies in *A. thaliana* (38,43), may already have been present in the perennial ancestor. During *FLC* dial-up, both species showed two transcriptional signatures as previously reported in *A.halleri* (20,33), indicating that these behaviours are conserved across species in the field. In spring, *FLC* and *VIN3* levels correlated inversely, a pattern not captured in laboratory studies where abrupt cold-to-warm shifts rapidly extinguish *VIN3* expression (24). Moreover, *VIN3* reached its seasonal minimum before *FLC* attained its maximum, raising the possibility that the absence of *VIN3* is a permissive condition for re-establishing high *FLC* levels. Finally, *FLC* dial-up kinetics differed between species and were mirrored in *FT* peak shapes: in *E. japonicum*, the rapid *FLC* dial-up coincided with a narrower *FT* peak, whereas in *A.halleri* the slower *FLC* dial-up corresponded to a broader *FT* peak. This could be consistent with a FT-FLC feedback loop evidenced in the winter annual life history (44,45).

The conservation of the VIN3–FLC–FT domino-like sequence between *E. japonicum* and *A.halleri* suggests that the roles of *AhgFLC* and *AhgFT* may not be as tightly linked to reversion and flowering transitions, unique to the life history of *A.halleri*, as previously proposed (18,33), as it was also recently reported (46). The mechanistic or biological basis within the VIN3–FLC–FT regulatory module awaits genetic analysis in their respective perennial life histories, which lie beyond the scope of this study. Instead, we regard the conserved features of the VIN3–FLC–FT network as the empirical basis for applying a more generic comparative measure—the thermal memory interval—introduced in the next and the last sections.

### Thermal memory interval adds predictive power to chromatin and expression features of spring and autumn *FLC* states

Previous work established that *AhgFLC* expression follows a seasonal cycle characterized by chromatin-based stability during both *AhgFLC* dial-down and *AhgFLC* dial-up, while these phases remain sensitive to the changing trend of temperatures in autumn and spring (21,23). First, we addressed whether this feature is conserved in the distantly related perennial *E. japonicum*.

In both species, *FLC* expression reaches mid-intermediate levels at two timepoints per year: once during *FLC* dial-down in early winter and again during *FLC* dial-up in mid-spring (Fig 2 B,E). We selected these matched expression timepoints to test their temperature sensitivity in *E. japonicum*. Transfer experiments from natural conditions to two contrasting constant temperatures (4 °C and 23 °C) showed that temperature sensitivity differs seasonally (Fig 3A). In winter, *EjFLCs* expression remained stable for four weeks following transfer to 23 °C, while in spring, the same transfer led to a gradual ∼10-fold increase. Conversely, exposure to 4 °C caused a rapid *EjFLCs* decline in winter but had no effect in spring until four weeks later. These results mirror what was previously observed in *A.halleri* (*18,21*), confirming that the *FLC* dial-up and *FLC* dial-down phases are distinct in their responsiveness to temperature, even when transcript levels are matched. This represents an epigenetic hallmark: the same DNA sequence can give rise to different transcriptional outcomes depending on prior history. The differential behavior of these dialling states prompted us to verify whether corresponding distinct chromatin dynamics are conserved in *E. japonicum*.

**Fig 3.**
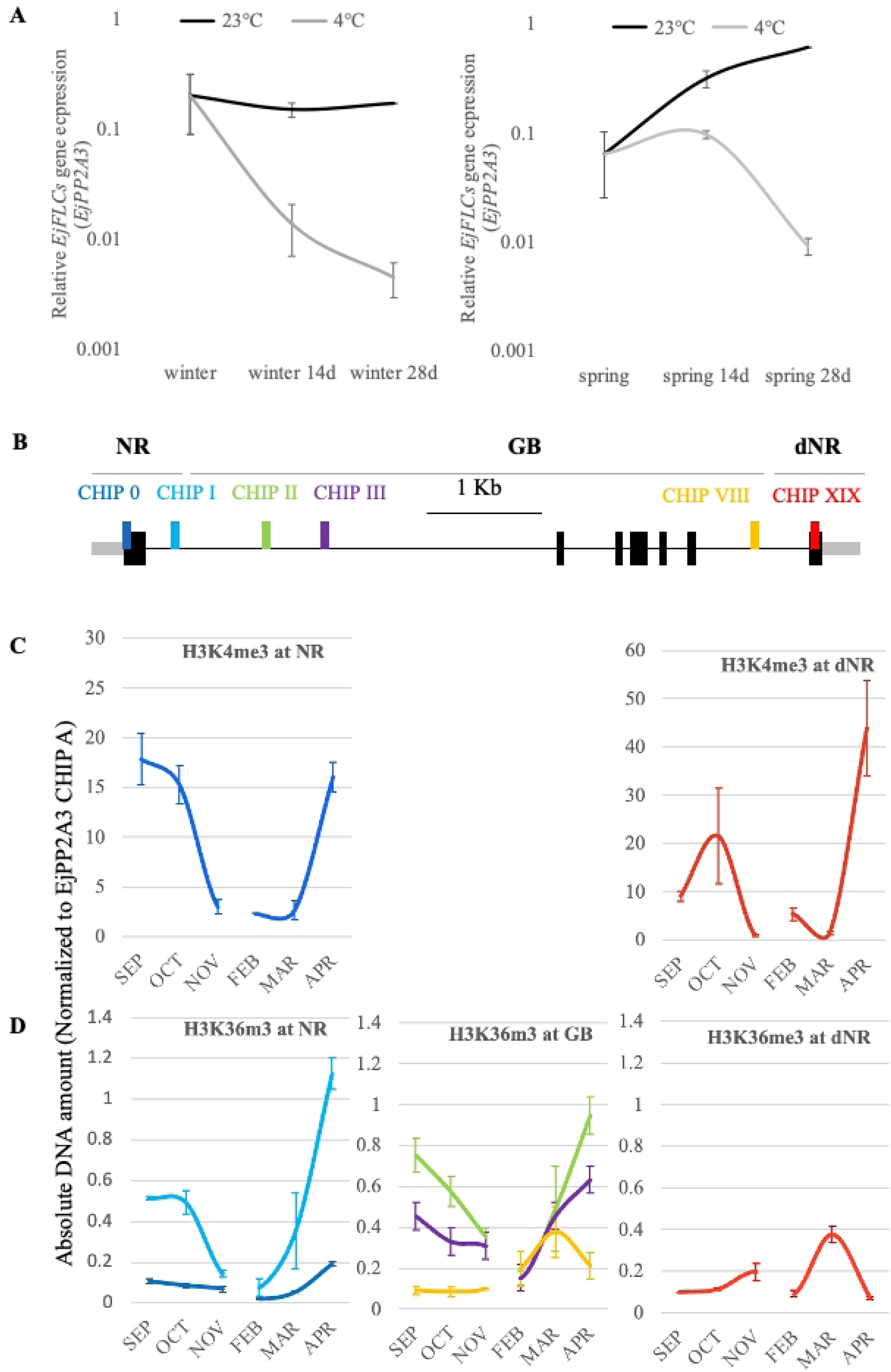
Asymmetry of *FLC* dialing states at gene expression and chromatin level in *E. japonicum.* (A) Distinct transcription states *EjFLCs* at onset of winter and spring. Average of three biological replicates of mRNA quantification of *EjFLCs*, as indicated on the vertical axis, using ΔΔCT method and *EjPP2A3* as an internal control. The scale for each gene was chosen to maintain consistency throughout the manuscript. Lines represent smooth curves. Plants were transferred from Ikawa on 17 December 2018, “winter”, and 18 March 2019, “spring”. (B) Distribution of ChIP amplicons relative to the *EjFLCs* with untranslated regions (grey boxes), introns (lines) and exons (black). CHIP 0 and I are in the Nucleation Region (“NR”), CHIP II, III, VIII in the gene body (“GB”) and CHIP IX in the distal Nucleation Region (“dNR”). The three chromatin domains are approximated by the grey lines on the top. The genomic features, but not the ChIP amplicons, are drawn to scale. (C) Absolute quantification of H3K4me3 at regions NR (left) and dNR (right), normalized to EjPP2A3 CHIP A. (D) Absolute quantification of H3K36me3 along all regions, normalized to EjPP2A3 CHIP A. Colour code is the same in B-D.

To address this, we mapped selected chromatin dynamics at *EjFLCs* (Fig 3B-D). We designed oligonucleotide probes targeting three regions previously defined in *AhgFLC*: the nucleation region (NR), gene body (GB), and distal nucleation region (dNR) (Fig 3B). Using ChIP-qPCR, we quantified active chromatin marks—H3K4me3 and H3K36me3—in the month proceeding the mid-intermediate FLC levels, during late autumn and late winter. At the NR, levels of H3K4me3 and H3K36me3 varied quantitatively with *EjFLCs* expression (ChIP 0, I) (Fig 5C), consistent with a role in modulating sense transcription. However, dynamics at other domains were distinct. At the GB, ChIP II reflected NR-like behavior, but ChIP III and VIII did not. These sites showed no decline in H3K36me3 from October to November (despite ∼15-fold transcriptional downregulation, Fig 5C) and no increase from March to April (during ∼30-fold transcriptional upregulation, Fig 5C). At the dNR (ChIP IX), H3K36me3 levels increased during autumnal dial-down—opposite to transcription—and decreased in spring during dial-up (Fig 3C, D, Fig 5C). These data indicate that, while the NR chromatin state aligns with expression, distal regions such as the GB and dNR maintain distinct histone modification patterns that are not transcription-coupled and vary across seasons. These distinct chromatin domains at *EjFLCs* match the organizational logic observed at *AhgFLC* (*21*) thereby supporting conservation in *E. japonicum*.

Next, we sought to distinguish the dialling states quantitatively by measuring the thermal memory interval of *FLC* expression specifically during the dialling phases —the time window of past temperature that best predicts mRNA levels—using simple moving averages (SMAs) and linear regression (Fig 4B; S4 Fig). During *FLC* dial-up (March–May), *FLC* integrated recent temperature changes over short intervals: 14–15 days in *E. japonicum* and 29–44 days in *A.halleri*. In contrast, during *FLC* dial-down (September–February in *E. japonicum*; November–February in *A.halleri*), memory intervals were substantially longer: 68–90 days and 57–107 days, respectively. This reveals a new feature of FLC regulation—season-dependent memory length—which is conserved across species. Importantly, the distinction in thermal memory intervals between *FLC* dial-up and *FLC* dial-down recapitulates mechanistic differences previously attributed to *cis*-based chromatin and *trans*-acting factors (23).Thus, TMIs provide a quantitative parameter of cellular memory length that aligns with known regulatory mechanisms but can be derived from expression and environmental data alone. In this way, TMIs extend mechanistic insights into a generalizable and comparative measure of memory across contexts.

**Fig 4.**
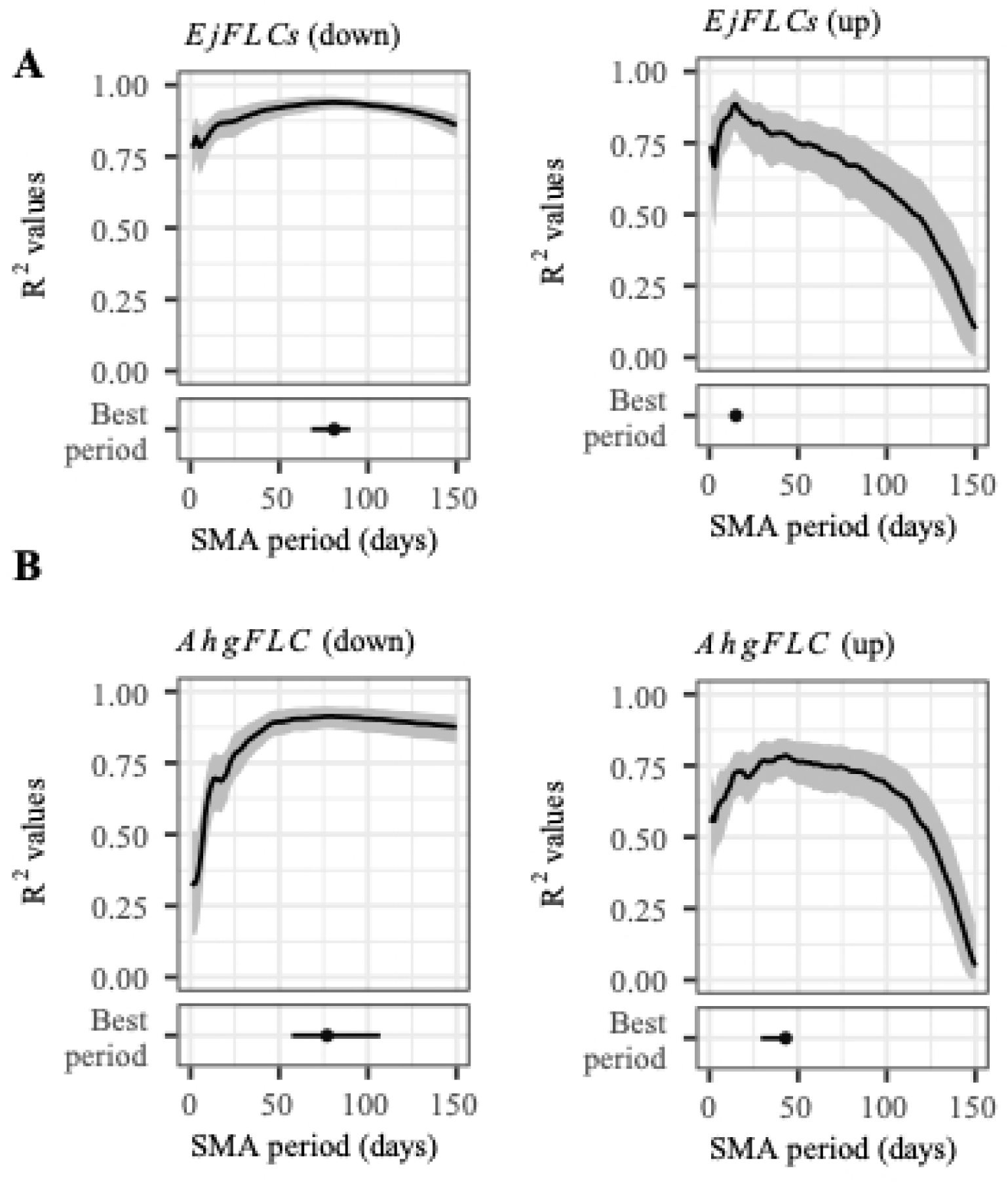
Linear regression analysis of EjFLCs (A) and AhgFLC (B) mRNA levels on the simple moving average of past temperature during dialing intervals. The results of linear regression analyses on the SMAs of the daily mean temperature with different window lengths specifically during *FLC* dial-up (right) and *FLC* dial-down (left) for *EjFLCs* and *AhgFLC* and R2 values are shown.

### VIN3–FLC–FT network maintains flexibility and robustness while encoding distributed memory

To examine how the VIN3–FLC–FT network responds to environmental conditions different from the native site and to assess the ability of this network to respond to long-past environmental histories, we conducted transfer experiments from the native site in Ikawa to a warmer location in Tsukuba (Fig 5A). Average hourly temperatures in Tsukuba were at least 4 °C higher than in Ikawa for most of the year (S8 Fig), a temperature difference previously shown to influence vernalization dynamics and flowering phenology in *A.halleri* (33). Flowering time and gene expression were monitored following transfers at the onset of *EjFLC* dial-down in September and October, over six to seven months. Flowering was delayed in the September transfer group and absent in the October transfer group (Fig 5A). These phenotypes may be consistent with elevated *EjFLCs* (both transfers) or reduced *EjFT* expression (October transfer) or both (September transfer), given that *EjFLCs* inhibit flowering (Fig 1E) whereas *EjFT* promotes it (31). Also, flowering phenotypes of our transfer experiments are consistent with warming studies in *A.halleri* (*47,48*).

**Fig 5.**
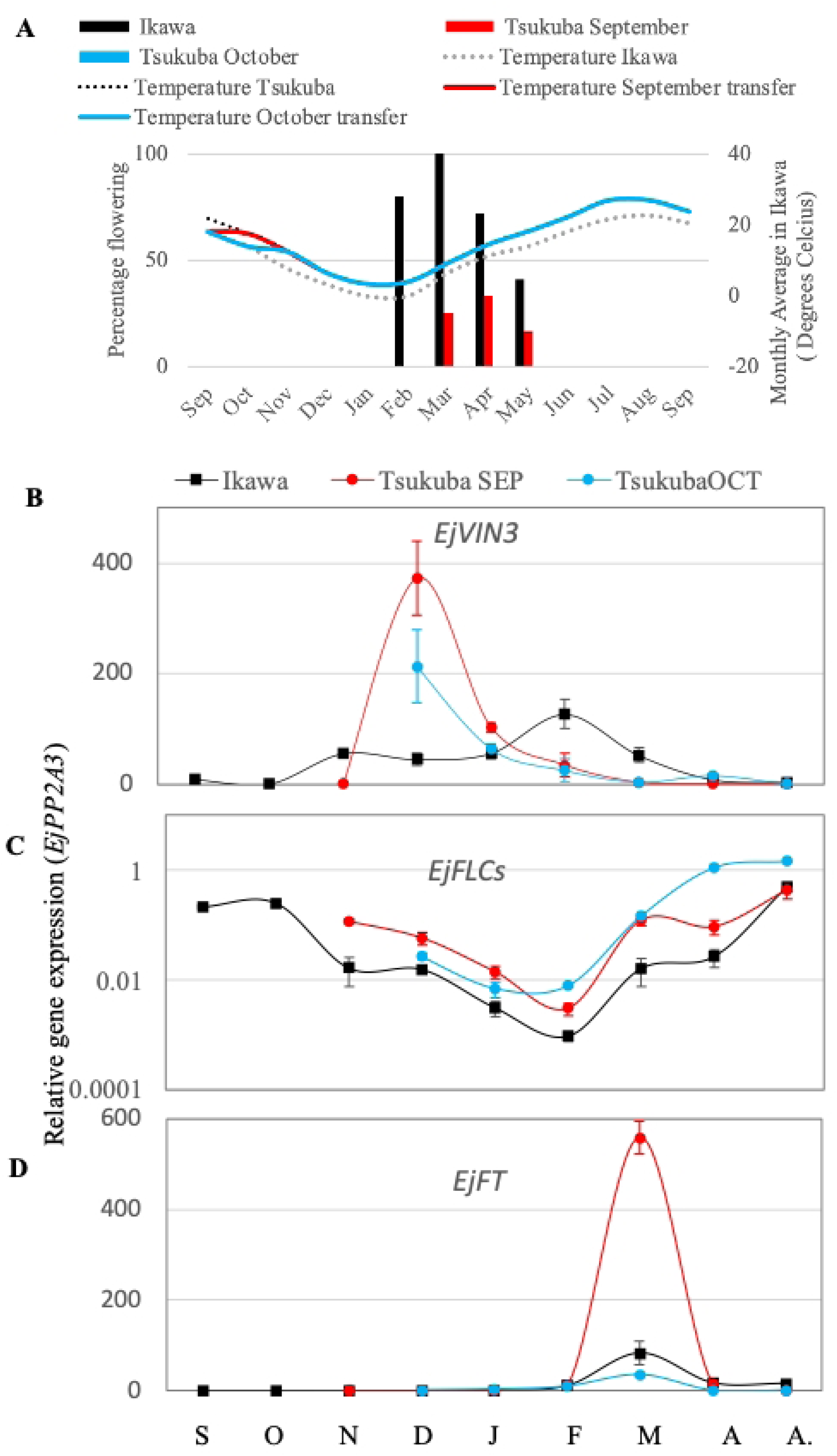
*E. japonicum* VIN3-FLC-FT module dynamics at the native and a remote site. (A) Flowering time against monthly average of temperature in transfer experiments from Ikawa to Tsukuba performed in September and October. (B-D) Gene expression analysis of *EjVIN3*, *EjFLCs* and *EjFT* in Tsukuba transfer experiments against the Ikawa control. Average of three biological replicates of mRNA quantification for all genes, as indicated on the vertical axis, using ΔΔCT method and *EjPP2A3* as an internal control. The scale for each gene was chosen to maintain consistency throughout the manuscript. Lines represent smooth curves. S, 28 September 2021; O, 26 October 2021; N, 25 November 2021; D, 28 December 2021; J, January 30 2022; F, 25 February 2022; M, 15 March 2022; A, 4 April 2022, A., April 2022.

Despite aberrant flowering, the VIN3–FLC–FT domino-like effect remained recognizable in the monitored cellular lineage, indicating that this regulatory sequence can operate beyond, and is not entirely coupled to, the floral transition. The characteristic four-quadrant expression pattern of *EjFLCs* remained recognizable (Fig 5B–D). Each gene displayed distinct response dynamics relative to the environment difference of the two transfers: *EjVIN3* diverged 2–3 months after transfer, whereas *EjFLC*s diverged throughout the entire interval analyzed. Strikingly, *EjFT* “memorized” the autumn environmental variation until spring, as indicated by the of change of amplitude of *EjFT* expression in March (Fig 5D). These observations indicate that individual network components can respond to environmental changes over different timescales while maintaining the overall regulatory framework, demonstrating the flexibility and robustness of the network. The differences in response timing among *EjVIN3*, *EjFLCs,* and *EjFT* suggest that environmental history is distributed across the network.

### Thermal memory intervals vary with genes and species and reveal distributed effects

While VIN3, FLC, FT genes showed broadly conserved expression patterns across divergent species (Fig 2), we sought to quantify these similarities more precisely, motivated by the need for a standardized metric and by our observation that memory timescales differ between genes and seasons (Fig 5B-D). We performed linear regression analyses relating gene expression to simple moving averages (SMAs) of daily mean temperature across multiple time windows (Fig 6; S5–S7 Figs). For each gene, the SMA window that best explained transcript abundance differed between genes and species (Fig 6). A consistent trend emerged within each species: *VIN3* integrated temperature over shorter windows (4–6 days for *EjVIN3*, 1–16 days for *AhgVIN3*), *FLC* over intermediate windows (13–28 days for *EjFLCs*, 43–48 days for *AhgFLC*), and *FT* over longer windows (104–150 days for *EjFTs*, 150 days for *AhgFT*). This indicates that thermal memory intervals are distributed along the regulatory cascade, possibly reflecting both each gene’s intrinsic regulatory features and contributions from upstream components. For example, *FLC*’s thermal memory interval, largely determined by its *cis* chromatin switch (8,10,21,23) could extend *FT*’s interval.

**Fig 6.**
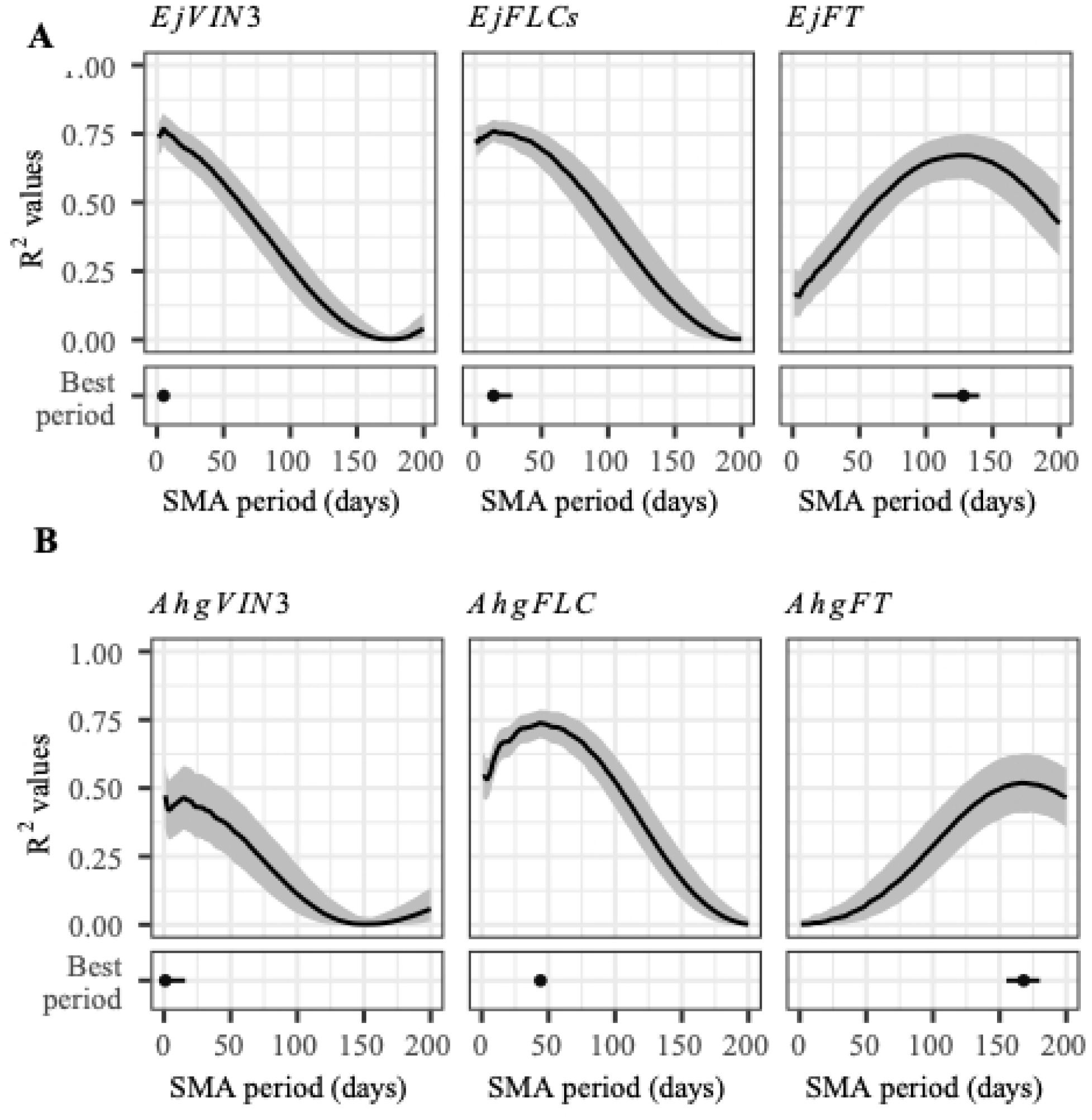
Linear regression of *VIN3*, *FLC*, and *FT* on the simple moving averages of past temperature in two perennials. The results of linear regression analyses on the SMAs of the daily mean temperature with different window lengths. R2 values for *EjVIN3*, *EjFLCs*, and *EjFT* (A) and *AhgVIN3*, *AhgFLC*, and *AhgFT* mRNA levels (B) are shown.

Our results provide the first estimates of thermal memory intervals for *VIN3* and *FT*, extending the concept beyond *FLC*. While these genes are also subject to chromatin regulation (49,50), their TMIs could be resolved without prior mechanistic knowledge, illustrating how the approach might generalize to other genes and contexts. FLC has long been recognized as a striking case of cellular memory, integrating past temperatures over ∼6 weeks (18). Here, we show that *FT* integrates temperature over even longer periods, up to ∼21 weeks, revealing that thermal memory can operate across broader timescales and highlighting the potential for TMIs to uncover similar patterns in other genes and species. The conservation of this progressive pattern across two divergent *Brassicaceae* suggests shared, modular principles for encoding seasonal environmental history independent of the perennial type of life history.

While TMIs provide a powerful and generalizable metric for cellular memory, their limitations should be acknowledged. Calculating TMIs requires dense, high-resolution gene expression and environmental data, such as the biweekly census collected over two years in this study. TMIs capture memory timescales but do not reveal underlying molecular mechanisms, including *cis*-encoded chromatin states, trans-acting regulation, or feedback loops. Long intervals—such as those observed for *FT*—may reflect cumulative contributions from upstream components rather than intrinsic memory at that gene. Nevertheless, TMIs offer a versatile, comparative framework for quantifying cellular memory across genes, regulatory hierarchies, and species. Moreover, high-resolution gene expression data can be analysed against additional variables—such as anatomical or physiological traits (51) or functional readouts such as photosynthetic efficiency (52)—extending the utility of TMIs to capture how genes integrate fluctuating signals beyond temperature.

## Conclusions

Comparative field-based studies of *A.halleri* and *E. japonicum* revealed that perennial *Brassicaceae* with divergent life histories nonetheless share core features of *FLC* and the VIN3–FLC–FT regulatory module. By relating gene expression to past temperature, we applied the TMI as a transferable, quantitative measure of cellular memory. Calculating TMIs required a high-resolution census of gene expression over two years under natural conditions, combined with meteorological data, enabling the capture of seasonal transitions, the distinction of autumn and spring *FLC* phases, and the quantification of distributed information flow across *VIN3*, *FLC*, and *FT*. Importantly, TMIs provide a unified readout of cellular memory without prior mechanistic knowledge. Because the TMI can be derived directly from high-resolution expression data alongside other dense datasets—such as environmental, anatomical, or physiological measurements—this approach is broadly generalizable beyond the three genes and single environmental variable examined here, offering a comparative framework for studying cellular memory across genes, species, and contexts.

## Acknowledgements

We acknowledge the help from the following people with initial setup of the Ikawa study: Kenichiro Hisada, Ryo Ohsawa, Chika Kasama, Kazuko Ito, Yasuko Shirai, Makiko Yamamoto, Lumi Matsuda. We thank Florin Bocaneala for initial advice on analysis of long-term data series and Jon Homewood for artwork in Fig 1.

## Author Contribution

DB designed and performed research, collected, analyzed data, interpreted data and wrote all the manuscript versions. HN performed research and analyzed data. OT analyzed data. YE, VE CF performed research. KY analyzed data; HK designed research. All authors read and commented on the manuscript versions.

## Supporting information captions

**Fig S1. Atypical perennial life history of *Arabidospis halleri gemmifera.*** Plants grown under “near-nature” conditions (windows were open to maintain near outdoor temperature) in a meshed glasshouse in Tsukuba from September 2023 to May 2024 were pressed at different stages of development. (A) Mature rosette (October) (B). Full flowering stage (early March). Note that the number of inflorescence stems (red arrows) is about the same with the number of rosette leaves. Since each rossette leaf is part of a repeating metamer composed of leaf, internode and subtended meristem, this implies that all vegetative meristems were induced to flower (C) Full reversion stage (early May), at the time all meristems, apical and axillary, have reverted into vegetative and developed into aerial rosettes (blue arrows).

**Fig S2. Sequence analysis of *FLC* coding regions in *Eutrema japonicum.*** (A) DNA sequence polymorphisms and resulting amino acid differences between *EjFLC1* and *EjFLC2.* Numbers indicate postion of the base pair from ATG codon. (B) Multiple sequence alignment between *AtFLC*, *EjFLC1*, *EjFLC2*. Amino acid differences between two species are in red. Arrow indicate amino acid difference between *EjFLC1* and *EjFLC2*.

**Fig S3. *EjFLCs* genes are not differentially expressed throughout the year.** Percentage of cDNA clones for each *EjFLC* gene, curated based presence of the six basepair substitutions from chromatographs from March 8, 2017, May 24, 2017, June 14, 2017, July 12, 2017, September 21, 2016, December 14 2016. The number of clones sequenced from each sample is indicted in brackets.

**Fig S4. Linear regression of *EjFLCs* (A, C) and *AhgFLC* (B, D) gene expression levels on simple moving average of past temperature with different window lengths during *FLC* dial up (A, C) and *FLC* dial down (B, D).** Gene expression level relative to *EjPP2A3* or *AhgPP2A3* was plotted against the SMA of past temperature with window length of 1day, 20, 40, 80 and 150 days for both genes. The time points used for the analyses can be found under “Statistical modelling” of the “Materials and Methods” section. Coefficient of determination (R^2^) is shown within the plots.

**Fig S5. Overlay between *FLC* expression and simple moving average of past temperature with different window lengths over 2 years in *Eutrema japonicum* (A) and *Arabidopsis halleri gemmifera* (B).** The simple moving average (SMA) of past temperature with window length of 1 day, 20, 40, 80 and 150 days during the two years are shown in different colors, together with *EjFLCs* (A) and *AhgFLC* (B) expression.

**Fig S6. Linear regression of *EjVIN3*, *EjFLCs*, *EjFT* expression levels on simple moving average of past temperature with different window lengths**. Gene expression level relative to *WjPP2A3* was plotted against the SMA of past temperature with window Length of 1day, 10, 20, 30, 40 days for *EjVIN3* (A) and *EjFLCs* (B) and of 1 day, 20, 40, 80, 150 days for *EjFT* (C). Coefficient of determination (R^2^) is shown within the plots.

**Fig S7. Linear regression of *AhgVIN3*, *AhgFLC*, *AhgFT* expression levels on simple moving average of past temperature with different window lengths.** Gene expression level (relative to *AhgPP2A3*) was plotted against the SMA of past temperature with window Lengthe of 1day, 10, 20, 30, 40 days for *AhgVIN3* (A) and *AhgFLC* (B) and of 1 day, 20, 40, 80, 150 days for *AhgFT* (C). Coefficient of determination (R^2^) is shown within the plots.

**Fig S8. Example of *E. japonicum* plants from Ikawa to Tsukuba transplant experiments.** (A) The twelve plants sampled in Ikawa on September 28 2021 on arrival in the lab in Tsukuba (one day after sampling in Ikawa) (B) and (C) Plants growing outdoor in Tsukuba, at the time of sampling in November and December.

**Fig S9. Location and annual hourly temperatures at the site of origin and remote site.** (A) Location of Ikawa and Tsukuba sites (B) Hourly temperatures in Ikawa and Tsukuba and the difference in temperature over interval 20.09.2021-20.08.2022.

**Table S1. *E. japonicum* sample repository.**

**Table S2. List of oligos use in this study.**

